# Helminth coinfection facilitates gammaherpesvirus infection in the wood mouse *Apodemus sylvaticus*

**DOI:** 10.64898/2026.05.11.723779

**Authors:** Kayleigh Newby-Gallagher, Jessica A. Hall, James P. Stewart, Parul Sharma, Simon A. Babayan, Amy B. Pedersen, Andy Fenton

## Abstract

Helminths are widespread parasites that can modulate host immunity, potentially increasing susceptibility to viral infections. However, evidence for these effects varies across systems and environments, and links between laboratory and wild populations remain unclear. We developed a tractable system using wood mice, *Heligmosomoides* spp. nematodes, and wood mouse herpes virus (WMHV) to bridge this gap. Combining laboratory and field experiments with population modelling, we examined how helminth infection, anthelmintic treatment and diet affect viral dynamics. Across lab and field data, helminth infection consistently increased WMHV risk, with stronger effects at higher worm burdens. Field results showed that anthelmintic treatment reduced viral infection, and laboratory experiments showed that improved nutrition mitigates helminth-induced increases in viral susceptibility. Our population-level modelling suggested that helminth burden-dependent facilitation can generate nonlinear effects on viral spread, dependent on helminth virulence. Our findings highlight the potential importance of helminths as facilitators of viral infections, and suggest that anthelmintic treatment may provide indirect benefits for viral control. We also show the value of integrating lab and field approaches on the same (or closely related) species, in particular the potential offered by the wood mouse – *Heligmosomoides* – WMHV system, to understand the drivers and consequences of host-helminth-viral interactions.

## Introduction

Helminths are ubiquitous parasites, causing chronic infections in many human and most wildlife and livestock populations. As such, these parasites form a common and persistent part of the host environment for any co-infecting microparasites, in particular co-circulating viruses. Helminths may be particularly important in viral coinfections since they can have immunosuppressive effects on the host [1-3], and skew the immune response towards a ‘Type 2’ (Th2) phenotype, downregulating the Type 1 (Th1) anti-viral response [4-9]. These changes to the host’s immune response can increase host susceptibility to and pathology caused by co-infecting viruses [9-12], and may shape the spread of viral diseases in animal and human populations [13-15]. Furthermore, if helminths facilitate viral infections, then we may expect an added benefit of anthelmintic treatment, through reduced susceptibility to viral infections [16].

However, there is conflicting empirical evidence about the impact of helminth coinfection, and of helminth removal via drug treatment, on co-infecting viruses [17-20]. Despite the potential for helminths to facilitate viral infections, studies in humans have found that helminth coinfection may decrease virus-specific immune responses [21], or lower viral loads [18] or have no effect on viral disease progression [22]. In wildlife, [23] found a positive association between the nematode *Heligmosomum mixtum* and Puumala Hantavirus infections (PUUV) in wild bank vole populations, whereas [20] found no significant association between nematode and Sin Nombre virus infection in *Peromyscus* spp. mice. Observations following anthelmintic treatments are similarly mixed. For example, Blish et al. (2010) [17] demonstrated that treatment targeting the nematode *Ascaris lumbricoides* in individuals coinfected with HIV-1 resulted in increased expression of CD4^+^ cells, and a trend for decreased HIV-1 RNA levels. However, other studies show no, or opposite, results of anthelmintic treatment on viral infections [18, 22]. For example, Modjarrad et al. (2005) [18] found no association between the treatment of helminth infections and HIV viral load in adults in Zambia. Additionally, Mulu et al. (2013) [7], studying HIV-1 infected individuals from Ethiopia found that anthelmintic treatment reduced HIV RNA levels in the plasma although, contrary to Blish et al. (2010) [17], there was no reduction in CD4^+^ cell counts. In a study of wildlife, Sweeny et al. (2020) [20] found that Sin Nombre virus infection levels in wild *Peromyscus* spp. mice increased after anthelmintic treatment. Together these studies show that helminth coinfection can alter many aspects of virus infection, host susceptibility and disease progression. However, those effects can be complex, potentially varying between host-helminth-virus combinations and environmental contexts.

Much of our mechanistic understanding of how co-infecting parasites interact come from laboratory models of infection. However, animals under controlled laboratory conditions have very different immune responses compared to wild animals. For example, specific pathogen free (SPF) laboratory mice (*Mus musculus*) have immune systems more closely related to those of neonatal humans than adult humans [24] and laboratory mice in general have relatively naïve immune systems, likely due to the lack of antigenic exposure [24-28]. Furthermore, laboratory mice typically receive a constant supply of food and water [26-28], whereas animals in the wild are likely to experience variation in the availability and quality of food resources. Host nutritional status is known to alter a host’s ability to mount an effective immune response, such that animals under nutritional stress (e.g., in the wild, or on low quality diets in the lab) may be less able to cope with parasitic infections [29-32].

Together these factors raise questions about whether helminth-viral coinfection interactions measured in laboratory mice are detectable in their natural host population [33].

To facilitate translation from lab to field, we ideally need a host-helminth-virus system that is amenable to carrying out controlled, mechanistic-focussed coinfection experiments in the lab, paired with the ability to conduct experimental treatments of the same host-helminth-virus system in its natural setting. Here we seek to develop such a system, presenting results from paired lab and field studies of the coinfection dynamics of a gastrointestinal nematode (*Heligmosomoides polygyrus*, or its sister taxon *H. bakeri*) and a virus (wood mouse herpes virus; WMHV, formally *Rhadinovirus muridgamma 7*), in their natural host, the wood mouse, *Apodemus sylvaticus*. Wood mice are abundant rodents throughout northern Europe, commonly infected with a wide range of parasites (e.g., helminths, viruses, bacteria and protozoa) and have proven over many years to be highly amenable to (re)capture, parasitological, virological and immunological assaying, and experimentation (including monitoring the effects of anthelmintic treatment [34, 35]) in their natural setting. Furthermore, we maintain a wild-derived, but now laboratory-reared colony of outbred wood mice, and have isolated parasites and pathogens from wild wood mice to conduct controlled infection/coinfection studies under standard laboratory conditions [32, 36].

Wood mouse herpes virus (WMHV) is a gammaherpesvirus belonging to the family *Orthoherpesviridae* and is most commonly found in wood mice and other murid rodents [37, 38]. WMHV is related to a number of human herpesviruses (e.g., Human herpesvirus-8 (HHV-8; also known as Kasposi’s sarcoma-associated herpesvirus or KSHV, formally *Rhadinovirus humangamma 8*), Epstein-Barr virus (EBV) and herpesvirus Saimiri (HVS-2) [39-41], and the laboratory strain, MHV-68, which is commonly used in laboratory mouse models of infection [33, 42-45]. Gammaherpesvirus infections in mice typically begin with an active, lytic phase, which involves productive viral replication in the lungs [46, 47]. Lytic virus replication in the lungs can be detected from 1 day post infection (dpi), with peak titres at 7 dpi, and is usually cleared from the lungs by 10 dpi (Figure 1B; [46-48]). From 3 dpi, the virus spreads from the lungs to the spleen, where latent virus is detectable from 6 dpi, peaking ∼14 dpi before declining to low stable levels by day 90 dpi, where it establishes lifelong infection [40, 46, 48, 49]. Very few viral genes are expressed during latency, allowing the virus to evade immune control, but reducing the likelihood of virus transmission to other individuals [40]. However, Open Reading Frame 73 (*ORF*73) is critical to maintaining latent viral infection [50, 51], making it a valuable marker of chronic viral infection. Latent virus can reactivate into the lytic phase, enabling it to infect new hosts [45], with the *rta* (Replication and Transcription Activator) gene, primarily encoded by Open Reading Frame 50 (*ORF50*), playing a central role in viral re-activation [44, 45, 52]. The lifecycle of rodent gammaherpesviruses has generally been investigated in laboratory mice (*M. musculus*), however Hughes et al. (2010) [33] found that progression of lytic MHV-68 infection in the lungs and latent infection in the spleens of wood mice (*A. sylvaticus*) is similar to that in laboratory mice.

**Figure 1.**
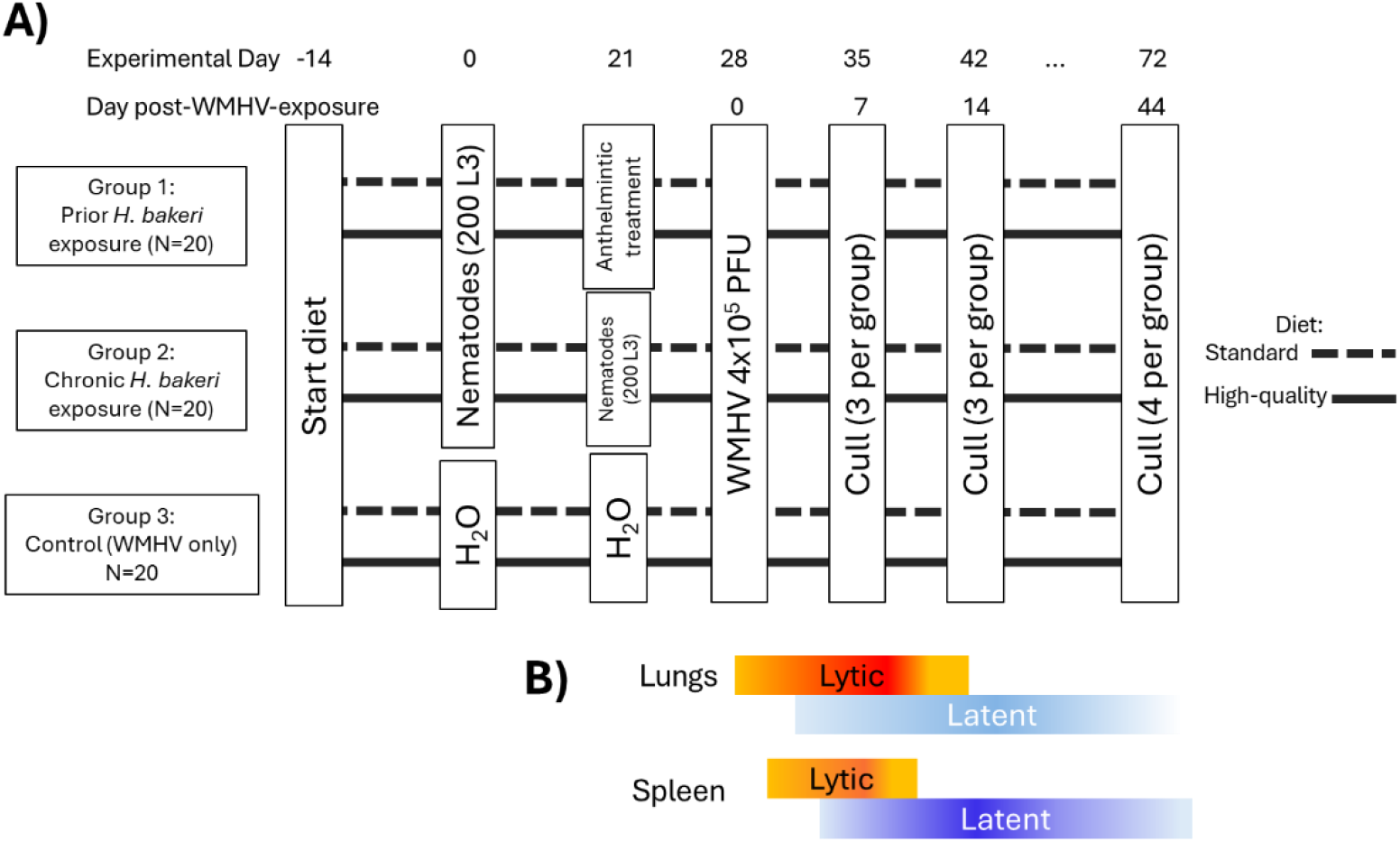
(A) Schematic diagram of lab experiment, and (B) summary of expected timings of lytic and latent WMHV infection in lungs and spleen, relative to ‘Days post-WMHV-exposure’ in the experimental design schematic; intensity of colours indicates intensity of viral loads. Summarised from [40, 46-49].

Epidemiological studies of wild wood mouse populations have found WMHV prevalence to be ∼10-30%, with infections tending to be more common in reproductively active males, indicating that male reproductive behaviours could provide an important route of transmission [37, 53, 54]. In addition, WMHV prevalence is highest in the spring, and adult mice infected with WMHV had recapture durations 25% shorter than uninfected mice, suggesting that infection may reduce the longevity of adult wood mice [37]. Wood mice are also typically infected with many other parasites, including helminths. The most common nematode infecting wood mice in the UK is *Heligmosomoides polygyrus*, which is highly prevalent (∼30-70%; [35]), and which is closely related to *H. bakeri*, known as a powerful immunomodulator, and a common laboratory model for studying helminth-immune system interactions [2, 55, 56]. Reese et al. (2014) [42] showed that infection of laboratory mice with the nematode *H. bakeri* induced expression of the *ORF50* gene in MHV-68, causing MHV-68 to reactivate from latency. Hence, we may expect the closely related species *H. polygyrus* to similarly increase WMHV infection in natural populations. Given the ubiquity of *H. polygyrus* in the wild, its potential for immune modulation, the high prevalence of WMHV in UK wood mice, the ability to use both parasites in lab infection experiments, and the general amenability of wood mice for lab and field studies, this system provides the ideal model with which to explore helminth-gammaherpesvirus coinfection interactions in both the lab and the field.

Here, we aimed to understand the relationship between helminth coinfection and WMHV infection dynamics in wood mice under both natural and laboratory conditions. Specifically, we asked (1) whether helminth (*H. polygyrus* in the field; *H. bakeri* in the lab) infection affects the probability or intensity of WMHV infection, (2) whether anthelmintic treatment alters WMHV infection dynamics and (3) under lab conditions, whether host diet moderates these effects. Finally, motivated by our findings we (4) develop a simple macroparasite-microparasite model to explore the population-level consequences of varying nematode burden and virulence on the basic reproduction number (*R*_*0*_) of WMHV, given the observed host-level effect of *H. polygyrus* on the probability of WMHV infection from our field data. We show that results from the lab and field correspond, with evidence that *H. polygyrus* infection facilitates WMHV infection risk and that the magnitude of this effect increases with worm burden. Furthermore, our field results suggest that reducing worm burdens through treatment can bring an additional beneficial effect of reducing the likelihood of WMHV infection. Our lab experiment further shows that host diet can play an important role in mediating the impact of helminth coinfection, such that animals on a high quality diet were able to mitigate the impact of helminth exposure on increasing the spread and duration of WMHV infection. Finally, our population model suggested that the burden-dependent facilitation of WMHV infection can result in nonlinear impacts of helminths on the basic reproduction number (*R*_*0*_) of the virus, dependent on the pathogenic impact of the helminths. Together, these findings highlight the potential role helminth coinfection can play in facilitating viral infections, at both the individual and population levels, and that helminth control via treatment can have indirect benefits for viral disease management.

## Methods

All field and laboratory protocols were carried out under Home Office Project Licence 70/8543 and with approval from the University of Edinburgh Animal, Welfare and Ethics review board (AWERB).

### Field experiment methods

Wood mice were collected from wild populations in Falkirk, Scotland from four field experiments carried out between 2013-2016 [35, 57, 58]. All small mammal trapping was conducted in mixed woodlands using Sherman live traps (H.B. Sherman 2 × 2.5 × 6.5-inch folding trap) baited with food (mixed grain and seeds, carrot/apple) and bedding, placed in pairs every 10m in 70 × 70m square trapping grids (see above citations for further details of field protocols).

Sampling designs differed between years. In 2013, we undertook cross-sectional sampling, in which wood mice were trapped over five nights in October and culled via a Schedule 1 method. We collected and stored spleens in RNA later at -80 °C for subsequent evaluation of WMHV infection status (details below). We also measured *H. polygyrus* infection burdens (numbers of adult worms) from the small intestine. The cross-sectional nature of this sampling resulted in contemporary measures of WMHV and *H. polygyrus* status at the time of capture for each individual mouse.

In 2014-2016, we employed longitudinal, capture-mark-recapture sampling where, at first capture, all wood mice were given a subcutaneous passive integrated transponder (PIT) tag, each with a unique 9-digit number, so they could be identified upon recapture. We sampled trapping grids for up to a month (2014: 14 days; 2015: 27 days; 2016: 30 days). To assess the effects of reduction of gastrointestinal nematodes (specifically *H. polygyrus*, the most prevalent species) on WHMV infection status, we randomly treated half of the individuals on each grid at first capture, with a safe and effective anthelmintic drug combination.Treated individuals were given a combined body weight-adjusted oral dose of Ivermectin at 100 mg/kg body weight, which targets adult stages of nematodes, and Pyrantel at 9.4 mg/kg, which targets larval/adult stages [59, 60], or water as a control. We have previously found that this combined drug treatment is very effective at reducing *H. polygyrus* infection for ∼10 -16 days [57].

Around ∼12-15 days after first capture, animals were humanely sacrificed. As in 2013, we collected spleens, which were stored in RNA later at -80°C, for evaluation of WMHV infection status, and analysed the gastrointestinal tract to quantify *H. polygyrus* worm burdens. These end-point samples allowed cross-sectional assessments of contemporary nematode–WMHV associations, consistent with the cross-sectional data from 2013.However, the additional longitudinal data of known individuals in the 2014-2016 experiments also allows us to test how removal of *H. polygyrus* through drug treatment influences subsequent WMHV infections.

### Lab experiment methods

Wood mice (*A. sylvaticus*) from our formerly-wild, but now lab-reared colony were maintained under standard laboratory conditions at the University of Edinburgh [32, 36]. The colony has been in captivity for many generations, but are purposely outbred to maintain genetic diversity. In this experiment, all mice were co-housed by sex within individually ventilated cages (IVCs; Techniplast, 1285L) with food and water *ab libitum*. At the start of the experiment, all mice were between 10-20 weeks old, and had not been exposed to *H*.*polygyrus/baker*i or WMHV.

Wood mouse herpes virus (WMHV) was isolated from wild wood mice [33, 61]. To test for the impact of nematode coinfection, we used the laboratory-passaged species *H. bakeri* from Prof. Amy Buck (University of Edinburgh), which is closely related to *H. polygyrus*, the species that naturally infects wood mice in the wild [55, 56], and which is well known for its ability to immunomodulate laboratory mice, modifying a wide range of host immune responses [62-64].

To test how nematode coinfection impacts WMHV infection dynamics in wood mice, we included three experimental groups (Figure 1A): 1) prior *H. bakeri* infection followed by subsequent WMHV infection, 2) chronic *H. bakeri* infection followed by WMHV infection, and 3) WMHV infection only (no *H. bakeri* infection). In each of the three groups, we had 20 age-and sex-matched wood mice. To test whether diet quality impacted WMHV and nematode infection dynamics, we randomly assigned 10 mice of each of the above three experimental groups to either a high-quality diet (‘Transbreed’ SDS; whole-diet nutrition with 20% protein, 10% fat, 38% starch, and high levels of micronutrients) or a standard maintenance diet (RM1, SDS; 14% protein, 2.7% fat, 45% starch, with standard levels of micronutrients; see [32] for further details). 14 days before the start of the experiment, all wood mice were co-housed by sex, with cages including individuals from all three experimental groups, and given their experimental diets. This resulted in 60 mice in total (3 experimental groups by 2 diets; 10 mice (5 females and 5 males) per diet-and-experimental group combination; Figure 1).

At Day 0, all mice from groups 1 and 2 received 150 *H. bakeri* larvae in 150μl of water via oral gavage. Mice in group 3 (WMHV only), received 150μl water via oral gavage. On Day 21, all mice in group 2 (chronic nematode infection) received a second dose of 150 L3 larvae in 150µl of water via oral gavage. All mice in group 1 (prior nematode exposure) received a combined weight adjusted dose of the anthelmintic Ivermectin and Pyrantel (9.4mg/kg Ivermectin and 100mg/kg Pyrantel) via oral gavage to clear their nematode infections. These treatments were intended to separate out the effects of on-going nematode infections on WMHV infections from prior nematode exposure. Group 3 (nematode control) mice received another 150µl of water via oral gavage. On Day 28, all mice were inoculated intranasally with 4×10^5^ plaque forming units of WMHV in 40μl PBS under light anaesthetic (isoflurane). Finally, on Days 35 and 42 and 72 (corresponding to 7, 14 and 44 days post WMHV infection, respectively), either three mice (for days 35 and 42) or four mice (for day 72) from each group were culled and lungs and spleens harvested and stored in RNAlater at -80 °C for subsequent quantification of WMHV infection status. These assays were timed to coincide with key phases in the infection cycle in the different tissues (Figure 1B).

### Quantification of WMHV infection status

From the field experiment, all available spleen samples for 2014 (n=15), 2015 (n=11) and 2016 (n=6) were used in RT-qPCR for WMHV detection. For 2013, 32 animals were selected to include an even number of male and female mice and *H. polygyrus* status (positive/negative). Any individuals with spleens that fell outside the interquartile range in spleen sizes for the population were excluded, as very large spleens could be the result of overactive spleens (splenomegaly) from other concomitant infections. Both spleens and lung tissues were used from the lab experiment. However, tissues from two mice (both on high quality diet from the control group that did not receive nematode larvae) were not able to be processed, leaving 58 spleen and lung samples for analysis.

WMHV infection status was quantified from each tissue sample using RT-qPCR. RNA extraction from spleens (2013) was carried out using RNeasy® PowerLyzer® Tissue & cells kit (Qiagen) according to instructions, including on-column DNase 1 treatment. Optical density of all RNA samples was measured to determine RNA concentration before running an agarose gel to ensure there was no DNA contamination. cDNA was reverse transcribed from RNA by first heating 2μg total RNA in the presence of random primers, dNTPs and RNase free water (total volume of 13μl) to 65 °C, before snap cooling on ice and adding first strand synthesis buffer plus 0.1M DTT, then heating to 42 °C before adding 1μl RTase (SuperScript II) and incubating at room temperature. The mixture was then incubated at 42 °C for 2 hours, then inactivated at 70 °C. Samples were stored at -20 °C until needed.

The purified cDNA was used to amplify WMHV *ORF50* (encoding the key lytic switch protein, Rta) and *ORF73* (latency associated) genes. Primers were designed for both *ORF50* and *ORF73* primers using the WMHV genome and tested on known WMHV-positive samples. Primers for PCR *ORF*50 (*ORF50*F: 5’-CATCTGAGGACGCGTTCATC-3’ and *ORF50*R: 5’-CAGTGACACATGTTCCAT-3’) generated a 100-bp product (Genbank Acc. No. AF105037.1), and *ORF73* (*ORF73*F: 5’ ACCATGCCAGGATCTATGYTATTGTGTGT-3’ and *ORF73*R: 5’-CTCCAAAGCATYTACTATTCAA-3’) generated a 157-bp product (Genbank Acc. No. GQ169129.1). A first PCR was carried out to check the validity of cDNA samples using the murine ribosomal protein L8 gene (Genbank Acc no. AF091511), using forward primer 5’-ACCAGAGCCGTTGTTGTTGTTGTGG-3’ and reverse 5’-AGTTCCTCTTTGCCTTGTACTTGTGG-3’. End point PCR was used to determine the presence of *ORF73* and *ORF50*, where the reaction mixture contained 5x PCR buffer, 10x dNTPS (2mM), Taq II Pol (0.8U/μl) and each primer at 10μM each. Additionally, *ORF*73 was analysed by RT-qPCR using the same reaction mixture as for PCR, but with the addition of 10x SYBR green 1. The PCR programme was as follows: 35 cycles of 2 min at 95 °C, 10s at 94 °C, 20s at 60 °C, 15s at 72 °C, 10s at 75 °C, followed by a melt curve analysis of 1 min at 94 °C, 30s at 65°C and then 30s at 95 °C.

### Quantification of H. polygyrus infection status

Adult worm burdens from the field experiment were quantified from culled animals by examination of the small intestine, caecum and colon of dissected animals, under a dissecting microscope, as described in [36, 57].

### Statistical analyses

We used generalised linear models (GLMs) and generalised linear mixed models (GLMMs) to explore the relationships between nematode-related drivers on measures of WMHV infection status in the field and lab experiments. We used three measures of WHMV infection status as the response variable in turn: (1) presence/absence (binomial models, based on positive expression of either *ORF*73 or *ORF*50); (2) the quantity of *ORF*73 gene expression, as a proxy for viral load, among positive samples (Gaussian models, using log(*ORF73* expression level) as the response variable); and (3) the probability of lytic infection among WMHV-positive samples (binomial models, based on expression of *ORF50*, among samples that were positive for either *ORF73* or *ORF50*).

For each of the above sets of analysis we explored the association of the relevant WMHV metric with each of several nematode-related explanatory variables. For the field data we explored: (1) end-point *H. polygyrus* presence/absence (binary factor), based on observed adult *H. polygyrus* infections at the final capture point of each animal, using data from all untreated animals in the four sampling years (2013-2016); (2) end-point *H. polygyrus* burden (adult worm count; continuous), again from untreated animals at their final capture point; and (3) lagged effects of previous treatment (factor: treated or not), using the longitudinal data from 2014-2016. For the lab experiment, initial analyses showed no differences in WMHV infection status between the two nematode-exposure groups (‘prior’, which were anthelmintic treated to remove infections, and ‘chronic’, which received two inoculations of *H. bakeri* larvae). We therefore combined them into a single ‘nematode exposed’ explanatory variable in the following analyses.

In addition to the focal nematode-related explanatory variables, we also included a fixed effect of host sex (factor: male or female) for analyses of both the field and lab data. For the field analyses we included Year (factor: 2013-2016) as a random effect to control for between-year differences, and for the lab analyses we included cull point (Factor: 1, 2 or 3, corresponding to days 7, 14 and 44 respectively post-WMHV exposure) as a fixed effect. For the lab experiment, we initially restricted analyses to just the data from the standard diet treatments, before exploring whether any nematode-related effects on WMHV status differed from those of the high-quality diet by re-running analyses for all the data, including an interaction between WMHV status and Diet (factor: standard or high-quality).

Hypothesis testing relating to the effects of the nematode-related variables on WMHV infection status was carried out by likelihood ratio tests (LRTs) comparing models with and without the relevant nematode-related variables. All models were inspected using standard diagnostic tools (residuals v fitted values, QQ plots, residual diagnostics using the DHARMa package [65]). All analyses were performed in R, v. 4.5.0 [66]. Mixed effect models were run using the *glmmTMB* package [67]; most GLM analyses were run using the ‘glm’ function, however the lab experiment data showed notable separation, so binomial GLMs of the lab data were run using the ‘brglm’ function from the *brglm* package [68].

### Population-level model: Predicting the effects of helminth burden and virulence on WMHV R_0_

Our field data analysis (see below) shows that increased *H. polygyrus* burden is associated with increased probability of being WMHV infected (Figure 2A). We used this finding to parameterise a simple epidemiological model to explore the population-level consequences of varying nematode burdens on the basic reproduction number (*R*_*0*_) of WMHV. Our model is based on the microparasite-macroparasite hybrid model of Fenton (2013) [69], extended to, and parameterised from, the ‘6-group SAL’ (Susceptible-Acute-Latent; male & female sub-groups) WMHV model of Erazo et al (2021) [54]. We emphasise this model is intended to explore helminth-gammaherpesvirus effects in general, and is not intended to accurately capture *H. polygyrus* – WMHV dynamics in wood mice specifically.

**Figure 2.**
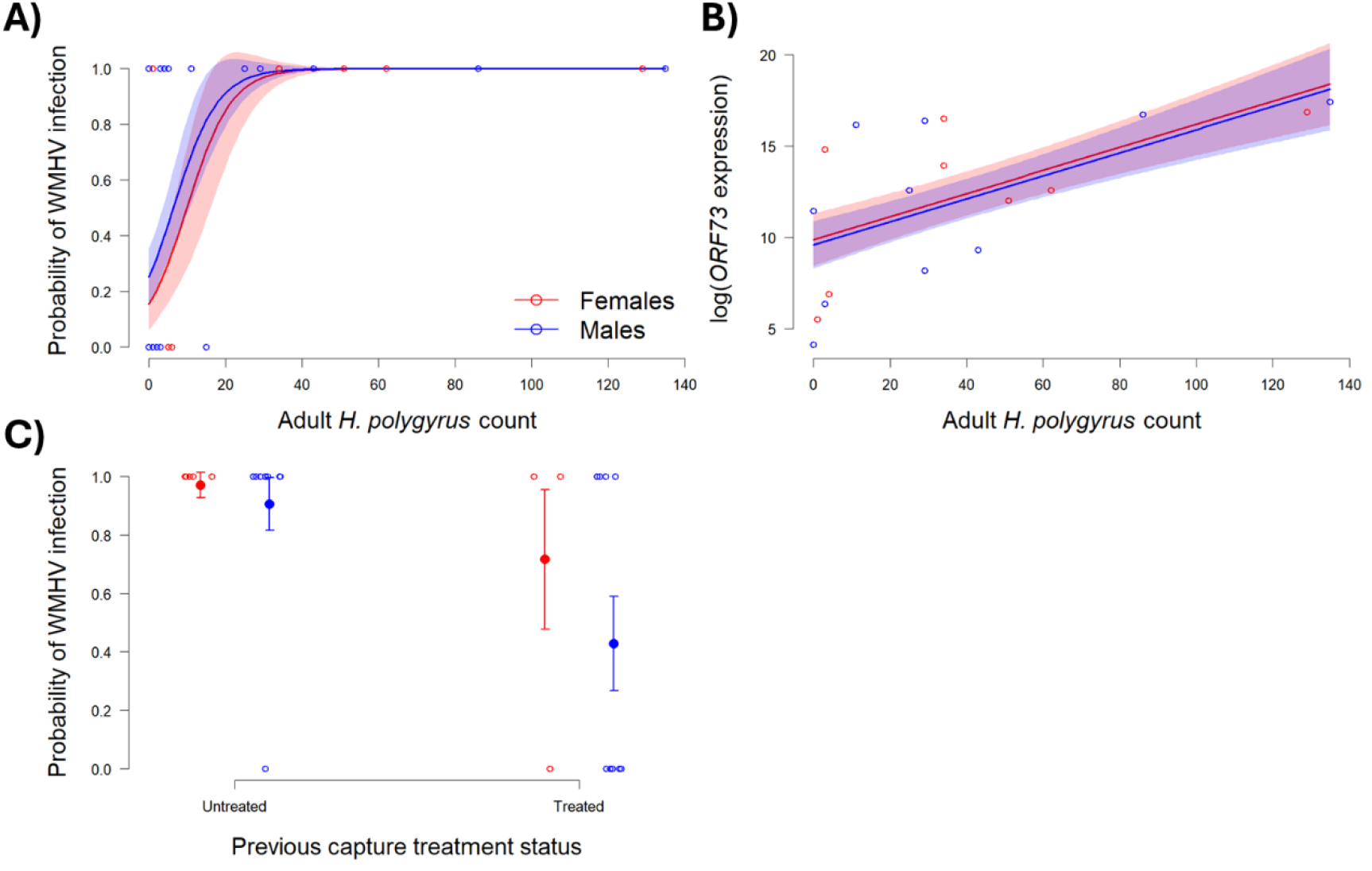
Field results. Relationships between end-point *H. polygyrus* adult worm count and (A) the predicted probability of WMHV infection and (B) log(*ORF73* expression), a proxy for viral load, and (C) relationship between prior anthelmintic treatment status and end-point WMHV infection status, each for females (red) and males (blue). Solid lines (for A and B) and points (for C) show mean predicted values; shaded areas (for A and B) and bars (for C) show ±1 SE. Open points show the raw data.

Erazo et al (2021) [54] showed that a 6-group model, incorporating density-dependent transmission within and between groups, best explained WMHV dynamics in a population of wood mice. This model tracked changes in the abundances of male and females in the population, broken down by WMHV-infection status in sex *i* (males, ‘*M*’, or females, ‘*F*’): Susceptible (*S*_*i*_), Active (infectious, *A*_*i*_) and Latent (non-infectious, *L*_*i*_) (Figure S1). Hence, the total number of individuals of sex *i* is *N*_*i*_ = *S*_*i*_ + *A*_*i*_ + *L*_*i*_ and the total number of WMHV-infected individuals of sex *i* is *I*_*i*_ = *A*_*i*_ + *L*_*i*_. In what follows we assume the total numbers of males and females (*N*_*M*_ and *N*_*F*_) are constant.

To incorporate the effect of *H. polygyrus* on WMHV transmission, we followed the approach of Fenton (2013) [69], and assumed the mean burdens of *H. polygyrus* in WMHV-infection class *x* of host sex *i, W*_*xi*_, are constant (i.e., we do not explicitly model helminth dynamics). These burdens were assumed to increase the mortality rate of each host class by per worm rate *α*. Hence the total dynamics of the system are described by the following set of equations (see Figure S1 for model schematic):

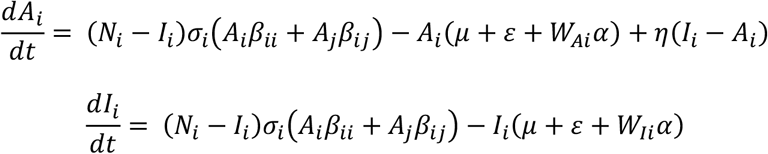

where: *σ*_*i*_ is the probability of infection of an individual of sex i given contact between a susceptible and infected host (effectively host susceptibility to WMHV infection); *β*_*ii*_ is the within-sex (male-to-male or female-to-female) contact rate; *β*_*ij*_ is the between-sex contact rate from individuals of sex *i* to sex *j*; *μ* is the background host mortality rate; *ε* is the transition rate from active to latent WMHV infections; η is the transition rate from latent to active hosts; and *α* is the per capita worm virulence (the increase in host mortality rate per infecting worm).

To incorporate the observed influence of *H. polygyrus* on WMHV transmission from our field data, we assumed the field-derived estimates of the effects of *H. polygyrus* burden on WMHV infection probability (Figure 2A) reflected burden-dependent changes in host susceptibility to the virus (*σ*_*i*_ for sex *i*). Hence, we used the sex-specific coefficients from our GLM analyses to generate the predicted probability of WMHV infection for each sex. All other parameters used the values estimated in Erazo et al (2021) [54] (see Supp Info, Table S1 for details).

In the Supp Info, we derive an expression for the basic reproduction number (*R*_*0*_) of WMHV using the next generation approach of Diekmann et al (2010) [70]. To explore the effect of changing *H. polygyrus* burdens on WMHV population-level transmission potential, we adjusted the observed worm burdens (*W*_*A*F_= 44.2; *W*_*AM*_ = 37.1) by a scalar, *W*_*adj*_, ranging from 0 (effectively eliminating *H. polygyrus* from the system) to 3 (trebling all mean worm burdens from the observed values), and measured the resulting effect on the derived expression for WMHV *R*_*0*_, which reflected changes in sex-specific susceptibility (*σ*_*i*_) and host mortality (via helminth *per capita* virulence, *α*). We repeated this for a range of hypothetical worm virulence values, ranging from *α* = 0 (completely benign) to *α* = 0.1 (highly pathogenic).

## Results

### Field experiment: summary statistics

A total of 58 spleen samples were analysed from wild wood mice for expression of viral *ORF*73 and *ORF*50 from 2013 – 2016. Across all years WMHV infection was detected (by either *ORF73* or *ORF50* positivity) in 47% (27/58) of samples (11 were both *ORF73-* and *ORF50*-positive; 12 were *ORF73*-positive but *ORF50*-negative; four were *ORF50*-positive but *ORF73*-negative). *H. polygyrus* was detected in 64% (37/58) of animals based on finding adult worms in the gastrointestinal tract; 34% (20/58) of animals were coinfected with both WMHV and *H. polygyrus*, and 24% of animals (14/58) were not infected by either.

### Field experiment: associations between nematode-related factors and WMHV

We found significant positive relationships between *H. polygyrus* burden (adult worm count) and both the probability of mice being infected with WMHV (likelihood ratio test (LRT) comparing models with and without *H. polygyrus* burden: χ^2^ = 10.914, df = 1, p = 0.001, model coefficient = 0.171; Figure 2A) and the quantitative level of *ORF73* gene expression (a proxy for viral load; LRT: χ^2^ = 7.596, df = 1, p = 0.006, model coefficient = 0.063; Figure 2B).On average, an increase of one adult worm was associated with ∼19% increase (Odds Ratio [95% CI] = 1.187 [1.008-1.400]) in the probability of being WMHV positive, and an increase of just over 1 unit in *ORF73* expression (Gaussian GLM coefficient (± SE) of ln(expression) =0.063 (± 0.0198)). We also found that animals which had previously received anthelmintic treatment were less likely to have detectable WMHV infection (by detection of either *ORF73* or *ORF50*) at their final capture compared to untreated animals (Figure 2C; LRT: χ^2^ = 6.102, df = 1, p = 0.014, model coefficient = -2.551; Odds Ratio [95% CI] = 0.078 [0.007-0.829]).

Given that most WMHV-positive samples were also *ORF73-positive* (12/27; 85%), the above results remained largely consistent if we used *ORF73* only as the indicator of WMHV infection status (Figure S2). There were no significant effects of any nematode-related variables (*H. polygyrus* presence/absence, burden or treatment) on the probability of lytic infection among WMHV-positive samples (expression of *ORF50* among samples that were positive for either *ORF73* or *ORF50*; LRT p > 0.05 for all analyses).

### Lab experiment: summary statistics

50 spleens (86% of the 58 samples tested) and 49 lung samples (84%) tested positive for WMHV infection via *ORF73* or *ORF50*. There was generally close correspondence between *ORF* markers: 75% of WMHV-positive spleen samples (37/50) were positive for both *ORF73* and *ORF50* (10 were *ORF73* positive only; three were *ORF50* positive only) and 80% of WMHV-positive lung samples (39/49) were positive for both (eight were *ORF73* positive only; two were *ORF50*-positive only). There was also consistency in WMHV infection status (by either *ORF* marker) between spleen and lung samples from the same animal: 72% (42/58) of animals were WMHV-positive in both tissues, seven were only positive in the lungs, eight were only positive in the spleen. One animal tested WMHV-negative in both tissues.

### Lab experiment: associations between nematode exposure and WMHV

Initial analyses showed no differences in WMHV infection status between the two nematode-exposure groups (‘prior’, which were anthelmintic treated to remove infections, and ‘chronic’, which received two separate inoculations of *H. bakeri* larvae). We therefore combined them into a single ‘nematode exposed’ variable in the following analyses. We also found no significant effects of nematode exposure on overall WMHV infection status (assessed by positivity of either *ORF73* or *ORF50*) in either spleens or lungs across the experiment, likely due to the very high levels of infection (∼85%) observed across all treatments.

We did find significant effects of nematode exposure on the probability of animals being *ORF73*-positive from both lung (LRT of brglm models: ΔDeviance = 6.922, df = 1, p = 0.009, model coefficient = 2.922; Figure 3A) and spleen (LRT: ΔDeviance = 6.334, df = 1, p = 0.012, model coefficient = 2.718; Figure 3B) samples. Animals exposed to nematodes had ∼19-fold increase in the probability of having an *ORF73*-positive lung sample (Odds Ratio = 18.58), and a ∼15-fold increase in the probability of having an *ORF73*-positive spleen sample (Odds Ratio = 15.14) compared to control mice. There were no significant effects of nematode exposure on the probability of lytic infection (*ORF50* positive among WMHV-positive samples) in either lungs or spleen.

**Figure 3.**
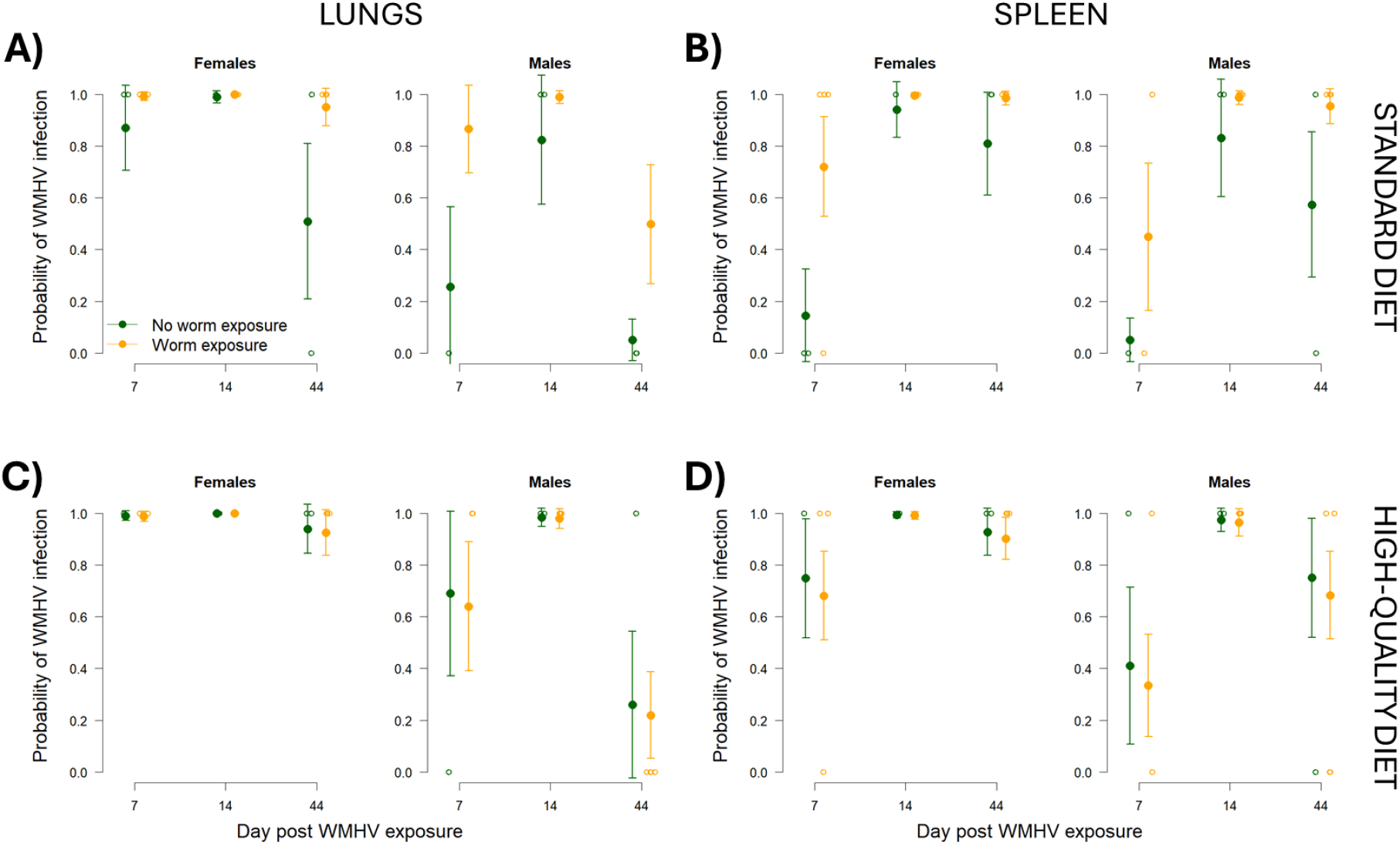
Lab results. Predicted probabilities of WMHV infection over days post exposure in lungs (left column) and spleens (right column) for females (left panel of each pair) and males (right panel of each pair) for animals previously exposed to *H. bakeri* (orange) or not (green).(A) and (B) for animals on the standard diet (RM1); (C) and (D) for animals on high-quality diet (Transbreed). Solid points show predicted means; error bars show ±1 SE. Open points show the raw data.

In the absence of nematode exposure (Figures 3A,B green points) we saw a general progression between days 7 and 14 post-infection of increased probability of detecting WMHV infections (by *ORF73*), initially higher in the lungs than the spleens. This peak was then followed by a reduction in infections by day 44 post-infection, again more dramatic in the lungs, as the infection progressed to the spleens. Among the nematode-exposed animals, there were less dramatic changes in WMHV infection levels over time, due to relatively high levels of sustained infection at both days 7 (early-phase) and 44 (late-phase) post-infection (Figures 3A,B orange points).

We found that mice on high quality diets differed from those on standard diets in the effect of nematode exposure on WMHV infections, although the effect in spleens was not significant at the ɑ = 0.05 level (LRTs comparing models with Nematode x Diet interaction to models without the interaction: Lungs: ΔDeviance = 4.737, df = 1, p = 0.030, Figure 3C;Spleens: ΔDeviance = 2.867, df = 1, p = 0.090, Figure 3D). In general, nematode-exposed and control mice on high quality diets did not differ in their WMHV infection risk (Figures 3C,D).

### Population-level model: Predicting the effects of helminth burden and virulence on WMHV R_0_

Our model predicted a unimodal relationship between worm burden and WMHV *R*_*0*_ (Figure 4). Increasing mean worm burdens from the observed values (*W*_*adj*_ > 1) was generally associated with a decline in WMHV *R*_*0*_, due to the progressively higher host mortality rate imposed by increasing worm burdens. The exception to this was when the worm was completely benign (*α* = 0; Figure 4 black line). Here there was no cost to the host (and therefore WMHV) of increasing worm burdens. However, there was very little additional benefit to WMHV, as the predicted impact of the observed worm burdens (at *W*_*adj*_ = 1) resulted in close to maximal susceptibility. Hence, further increases in mean worm burdens resulted in negligible increases in WMHV *R*_*0*_.

**Figure 4.**
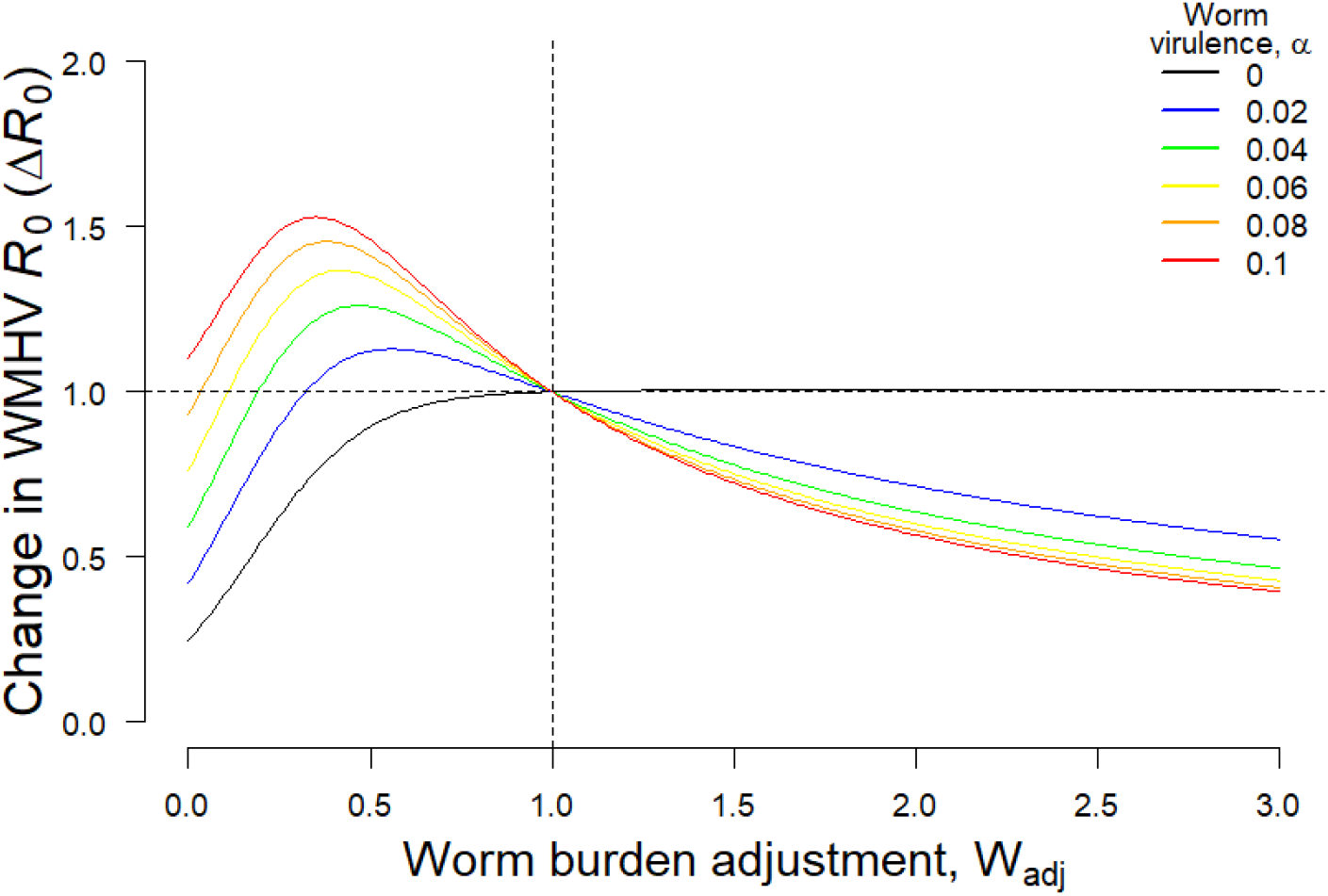
Model predictions of change in the basic reproduction number of WMHV (Δ*R*_*0*_), relative to the value at the observed mean worm burdens from the field data, due to changes in mean worm burden values (via the multiplier *W*_*adj*_), for differing hypothetical *per capita* worm virulence values (*α*). The vertical dashed line (*W*_*adj*_ = 1) corresponds to the observed mean worm burdens from the field data; the horizontal dashed line (Δ*R*_*0*_ = 1) shows the corresponding baseline at the observed mean worm burden.

Reducing mean worm burdens below the observed values by up to ∼50% generally resulted in an increase in WMHV *R*_*0*_, as the virus benefitted from the reduced cost of worm-induced host mortality, while retaining high infection probabilities at intermediate worm burdens. This effect was magnified for highly virulent worms (i.e., the benefit to WMHV of reduced worm burdens was greater as the worm was increasingly virulent). Again, the exception to this was when the worm was completely benign (Figure 4 black line), as there was no benefit to worm reduction, but a cost to WMHV from reduced worm-induced host susceptibility. Even for the more virulent worm scenarios, near-elimination of the worm populations (*W*_*adj*_ → 0) dramatically reduced WMHV *R*_*0*_, as the very low worm burdens were associated with very low probabilities of WMHV infection, bringing about a net reduction to WMHV, despite the alleviation of worm-induced host mortality. These unimodal relationships between worm burden and WMHV *R*_*0*_ are qualitatively similar to that predicted by Fenton (2013) [69] for a generic macroparasite-microparasite model with a positive interaction of the macroparasite on host susceptibility to the microparasite.

## Discussion

We show, through both lab and field experiments, that coinfection by a gastrointestinal nematode facilitates wood mouse herpes virus (WMHV) infections in wood mice.Furthermore, our lab experiment showed that the facilitative effects of helminths on WMHV infection risk can be mitigated through high-quality diets, potentially allowing better-resourced hosts to mount more effective anti-viral responses than animals on a standard diet. In addition, our field experiment showed that WMHV infection risk can be reduced by prior anthelmintic treatment, suggesting a possible added benefit of anthelmintic treatment programmes for coinfected individuals. However, our mathematical modelling showed that the consequences of reducing worm burdens on the potential for WMHV transmission (*R*_*0*_) can be complex, potentially resulting in increased viral transmission following helminth reduction, depending on the balance between helminth burden-dependent viral facilitation and host mortality. Together, these results show the value of combining lab and field studies of the same (or closely related) host-helminth-virus species, to better understand infection and coinfection dynamics, with relevance for wildlife, livestock and human helminth-viral systems.

Mechanistically, helminths can alter viral infection dynamics in several ways. Helminths generally stimulate T-helper 2 (Th2) immune responses, which are known to be antagonistic to anti-viral Th1 response [71, 72]. Furthermore, many nematodes, including those of the genus *Heligmosomoides*, are known to be powerful immunomodulators; indeed, the lab-adapted species *H. bakeri* used in our experiments is a model species to study immune modulation [2]. Hence hosts that are currently or have recently been infected with helminths may have reduced capacity to mount effective resistance against viral infections, which would generate results consistent with our findings of a facilitative effect of *Heligmosomoides* spp. on WMHV in both the lab and the field. In the specific case of gammaherpesviruses, which can transition between active (lytic) and inactive (latent) states, previous experimental work has shown that coinfection with *H. bakeri* induced expression of the *ORF50* gene in MHV-68 (related to WMHV), causing the virus to reactivate from latency to the lytic form [42]. We were unable to detect evidence of such an effect in either our lab or field studies, potentially due to limiting sample sizes or the temporal resolution of sampling, particularly in the field. Alternatively or additionally, Reese et al (2014) [42] conducted their experiment on MHV-68 in laboratory mice, a combination which may result in different outcomes from WMHV in wood mice. However, Hughes et al. (2010) [33] found the dynamics of MHV-68 infection in the lungs and spleens of wood mice was similar to that in laboratory mice. If the findings of Reese et al (2014) [42] do apply to WMHV in wood mice in the field, even though we failed to detect such an effect, this may result in *H. polygyrus* further increasing WMHV transmission via increased duration of the lytic form in coinfected individuals.

Our laboratory experiment found that mice on high quality diets exhibited no effect of helminth coinfection on WMHV infection dynamics. Hence, mice on high-quality diets were better able to cope with the dual challenges of helminth and WMHV exposure. Host nutritional status is well known to alter a host’s ability to mount an effective immune response against infection, with animals on low quality diets often shown to be less able to cope with parasitic infections [29, 30]. For example, Ezenwa (2004) [31] found that wild bovids unable to maintain adequate nutrition were less able to cope with gastrointestinal parasites, with reduced protein intake associated with lower resilience and resistance to infection. Furthermore, energy deficits in lab mice have been shown to simultaneously suppress both Th1 (INF-g) and Th2 (IL-4 and IL-5) responses and their effector functions [73], and IL-4 was also found to be suppressed in the spleens of zinc-deficient mice [74]. In our wood mouse system, we previously found in both wild and laboratory experiments that access to high-quality diet significantly increased resistance to *H. polygyrus* infection, enhanced anthelmintic treatment efficacy, and increased general and parasite-specific antibody responses [32]. While we were not able to assess worm burdens prior to WMHV inoculation in our laboratory experiment (as this would require culling mice), it is likely that mice fed the high quality diet had both lower worm burdens and/or a higher capacity to mount an antiviral response. While more experiments are needed to understand the mechanisms driving these results, our findings clearly show an impact of enhanced nutrition in potentially altering helminth-virus coinfection dynamics.

Motivated by our individual-level lab and field results, we modified an existing microparasite-macroparasite coinfection model [69] to explore potential population-level consequences of helminth coinfection increasing susceptibility to WMHV infection, via changes to the virus’ basic reproduction number, *R*_*0*_. Our model showed that viral *R*_*0*_ under such coinfection facilitation varies nonlinearly with helminth burden, dependent on the balance between helminth burden-dependent effects on host susceptibility to the virus and host mortality. High worm burdens generally result in a reduction in viral *R*_*0*_, due to high levels of helminth-induced host mortality reducing viral infection duration. However, reducing mean helminth burdens, for example through anthelmintic treatment, can increase viral *R*_*0*_ due to reductions in helminth-induced host mortality, potentially increasing the spread of the virus through the host population. The magnitude of this effect is strongest for highly virulent helminths. However, suppressing worm burdens further, ultimately to elimination, reduces viral *R*_*0*_, as the facilitative effect of helminths on host susceptibility to the virus is removed. These findings match previous studies [69, 75, 76] which show that the scaling of coinfection interactions between the individual and population scales can be counterintuitive. Of applied relevance, many helminths of human concern globally are targeted through Mass Drug Administration (MDA) programmes. Our experimental results suggest that if those helminths facilitate viral infections through increases in host susceptibility, then mass administration of anthelmintic drugs can potentially achieve an added benefit of reducing risk of viral infection. However, the consequences for the population-level spread of the virus can be complex, depending on the balance between helminth burden effects on viral facilitation and host mortality.

Overall, we show that the GI nematode *H. polygyrus*/*bakeri* facilitates wood mouse herpes virus infections in wood mice. Gammaherpesviruses like WMHV are highly common in humans - between 30%-90% of the world’s adult population is infected with at least one species of herpes virus, and a wide range of other animal taxa are similarly infected with gammaherpesviruses [77]. Given the relatively high prevalence of both herpes virus and helminth infections it is not surprising that coinfection between them is common, both in humans and in wildlife [37, 42]. Given the powerful immunomodulatory potential of many parasitic nematodes, it is likely that similar examples of viral facilitation by co-infecting nematodes may occur in a wide range of systems. The *Heligmosomoides* – WMHV – wood mouse system provides the capacity to directly translate between controlled coinfection experiments in the lab and the increased complexities of the natural setting, providing deeper insight into the causes and consequences of helminth-virus coinfection dynamics by lab or field studies alone.

## Acknowledgements

This work was funded by a NERC PhD studentship via the ‘*Adapting to the Challenges of a Changing Environment*’ DTP (NE/S00713X/1) to K. N-G, a NERC grant to A.B.P and A.F (NE/R011397/1), a Wellcome Trust Strategic Grant for the Centre for Immunity Infection and Evolution (095831) to A.B.P., a University of Edinburgh Chancellors Fellowship to A.B.P., as well as Wellcome Trust Institutional Strategic Support Fund (ISSF) grants to A.B.P. (ISSF 2014; J22737) and S.A.B. (097821/Z/11/Z). We are very grateful to Melanie Clerc, Amy Sweeny, Rowan Bancroft, Saudamini Venkatesan and many others for fieldwork and sample collection. We thank Bahram Ebrahimi for advice and experimental design help associated with measuring WMHV infection dynamics. We also thank Amy Buck and Elaine Robertson for providing the *H. bakeri* larvae for our laboratory experiment. We thank the Forestry Commission for permissions for fieldwork in Callendar Park.

All data and code scripts associated with this work are available in the Supplementary Material associated with this paper.

We have no competing interests.

