## Supplementary Information for "Helminth coinfection facilitates gammaherpesvirus infection in the wood mouse *Apodemus sylvaticus*"

*Calculating the basic reproduction number (R_0_) of WMHV*

To quantify the effect of *H. polygyrus* on WMHV transmission, we derived an expression for the basic reproduction number (*R_0_*) of WMHV using the next generation approach of Diekmann et al (2010) [1]. We focus on the generation of active (lytic) infections as these are the source of new infections (i.e., at the point of introduction of WMHV we assume all infections are initially lytic). Based on the above equations, we generated expressions for the ‘transmission’ matrix (*T*) and ‘transition’ matrix (*G*):

$$T= \left| \begin{matrix} N_{M}\beta_{MM}\sigma_{M} & N_{M}\beta_{MF}\sigma_{M} \\ N_{F}\beta_{FM}\sigma_{F} & N_{F}\beta_{FF}\sigma_{F} \end{matrix} \right|$$

$G= \left| \begin{matrix} -\left( W_{AM}\alpha+\mu+\varepsilon+\eta\right) & 0 \\ 0 & -\left( W_{AF}\alpha+\mu+\varepsilon+\eta\right) \end{matrix} \right|$,

resulting in the next generation matrix *K* = -*TG^-1^*:

$$K= \left| \begin{matrix} \frac{k_{11}}{D_{M}} & \frac{k_{12}}{D_{F}} \\ \frac{k_{21}}{D_{M}} & \frac{k_{22}}{D_{F}} \end{matrix} \right|.$$

where $k_{11}= N_{M}\beta_{MM}\sigma_{M}$, $k_{12}= N_{M}\beta_{MF}\sigma_{M}$, $k_{21}= N_{F}\beta_{FM}\sigma_{F}$, $k_{22}= N_{F}\beta_{FF}\sigma_{F}$,

$D_{M}= W_{AM}\alpha+\mu+\varepsilon+\eta$ and $D_{F}= W_{AF}\alpha+\mu+\varepsilon+\eta$.

*R_0_* is given by the dominant eigenvalue of *K*:

$$R_{0}= \frac{D_{F}k_{11}+D_{M}k_{22}+ \sqrt{\left( D_{F}k_{11}-D_{M}k_{22} \right)^{2}+4D_{F}k_{12}D_{M}k_{21}}}{2D_{F}D_{M}} .$$

To incorporate the observed influence of *H. polygyrus* on WMHV transmission from our field data, we assumed the field-derived estimates of the effects of *H. polygyrus* burden on WMHV infection probability (Figure 2A) reflected burden-dependent changes in host susceptibility to the virus ($\sigma_{i}$ for sex *i*). Specifically, we allowed the probabilities of WMHV infection given contact (the $\sigma_{i}$) to vary with mean worm burden by using the sex-specific coefficients from our GLM analyses (Figure 2A) to generate the predicted probability of WMHV infection for each sex (i.e., $\sigma_{i}$ becomes a function of mean worm burden of sex *i*: $\sigma_{i}=f(W_{Ai})$). At the observed mean worm burdens among male and female mice with active WMHV infections from our field data ($W_{AF}$ = 44.2; $W_{AM}$ = 37.1), the predicted infection probabilities from our GLM analysis were $\sigma_{F}$ = 0.997 and $\sigma_{M}$ = 0.995. All other parameter values relating to WMHV transmission were as estimated in Erazo et al (2021) [2] (Table S1).

We note that the previously-estimated net transmission rates (the $\beta_{ij}$ terms from Erazo et al (2021) [2]) are composite terms, implicitly comprising the *per capita* contact rate between host groups *i* and *j* $\left( \beta_{ij}^{C} \right)$, multiplied by the susceptibility of the ‘recipient’ host *i* ($\sigma_{i}$). i.e., $\beta_{ij}= \beta_{ij}^{C}\sigma_{i}$. Since we are independently calculating the susceptibility values of males and females (as described above), we derive the contact rate component of transmission between groups *i* and *j* as $\beta_{ij}^{C}= \frac{\beta_{ij}}{\sigma_{i}}$, where $\sigma_{i}$ is the susceptibility value calculated at the observed mean worm burden for group *i* from our data. We use these values of $\beta_{ij}^{C}$ as the (fixed) contact rates in our analyses, multiplied by the (varying) burden-dependent susceptibility value ($\sigma_{i}$) for sex *i*.

**Table S1**. Parameter definitions and values

| Parameter | Definition | Value* |
| --- | --- | --- |
| $S_{i}$ | Abundance of WMHV-uninfected mice of sex *i* | - |
| $A_{i}$ | Abundance of WMHV-acute (lytic) infected mice of sex *i* | - |
| $L_{i}$ | Abundance of WMHV-latent infected mice of sex *i* | - |
| $I_{i}$ | Abundance of WMHV-infected mice of sex *i* (acute + latent) | - |
| $N_{i}$ | Total density of mice of sex *i* (susceptible + acute-infected + latent-infected) | - |
| $\sigma_{i}$ | Susceptibility to WMHV (probability of WMHV infection given contact) for sex *i* | Dependent on mean worm burden^†^ |
| $\beta_{FF}$ | Female-to-female WMHV transmission rate | 0.0017 |
| $\beta_{MM}$ | Male-to-Male WMHV transmission rate | 0.0079 |
| $\beta_{MF}$ | Female-to-Male WMHV transmission rate | 0.0009 |
| $\beta_{FM}$ | Male-to-female WMHV transmission rate | 0.0032 |
| $\beta_{FF}^{C}$ | Female-to-female WMHV contact rate | $\beta_{FF}/\sigma_{F}$^**^ |
| $\beta_{MM}^{C}$ | Male-to-Male WMHV contact rate | $\beta_{MM}/\sigma_{M}$^**^ |
| $\beta_{MF}^{C}$ | Female-to-Male WMHV contact rate | $\beta_{MF}/\sigma_{M}$^**^ |
| $\beta_{FM}^{C}$ | Male-to-female WMHV contact rate | $\beta_{FM}/\sigma_{F}$^**^ |
| $\alpha$ | Helminth *per capita* virulence (additional host mortality rate) | varied |
| $\mu$ | Baseline host mortality rate | 0.060 |
| $\varepsilon$ | Rate of transition from acute to latent infection | 0.624 |
| $\eta$ | Rate of transition from latent to acute infection | 0.403 |
| $W_{AF}$ | Mean worm burden in active-infected females | 44.2 |
| $W_{AM}$ | Mean worm burden in active-infected males | 37.1 |

* units of rates are week^-1^

^†^ relationship between sex-specific susceptibility and mean worm burden derived from coefficients of binomial GLM: ‘WMHV infection probability ~ Worm burden + Sex’ (Figure 2A).

^**^ Using baseline susceptibility values calculated at the observed mean worm burden for sex *i*.

**Supplementary figures**

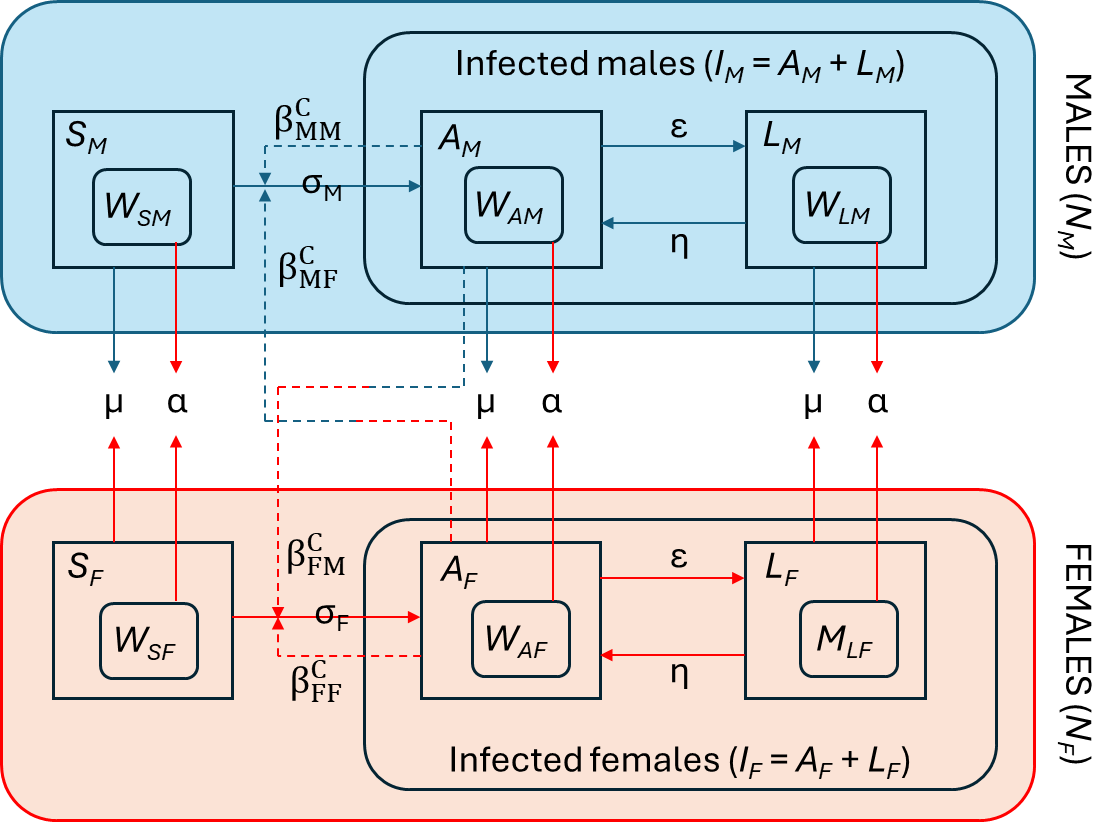

**Figure S1**. Schematic diagram of the population-level *R_0_* model, based on the 6-group (male-female; Susceptible (S) – Active (A) – Latent (L)) model of Erazo et al (2021) [2]. Parameters are defined in Table S1.

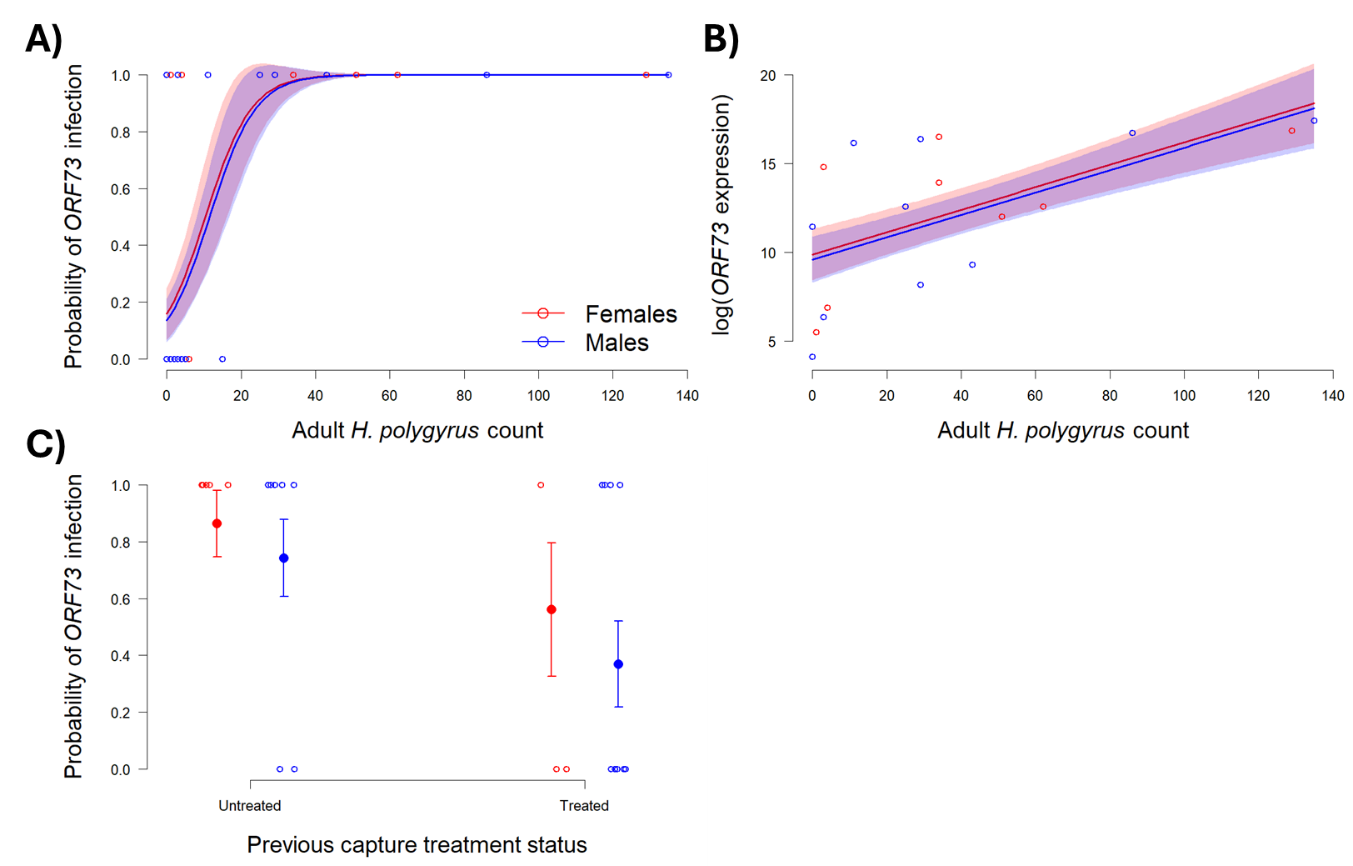

**Figure S2**. As in Figure 2 in the main paper, but showing results for *ORF73*, rather than combined WMHV status (infection determined by positive assays for either *ORF73* or *ORF50*). Relationships between end-point *H. polygyrus* adult worm count and (A) the predicted probability of WMHV infection and (B) log(*ORF73* expression), and (C) relationship between prior anthelmintic treatment status and end-point WMHV infection status, each for females (red) and males (blue). Solid lines (for A and B) and points (for C) show mean predicted values; shaded areas (for A and B) and bars (for C) show ±1 SE. Open points show the raw data.
